# The effect of parental age on the quantity and quality of offspring in *Syngnathus typhle*, a species with male pregnancy

**DOI:** 10.1101/2023.06.12.544574

**Authors:** Freya Adele Pappert, Daniel Kolbe, Arseny Dubin, Olivia Roth

**Affiliations:** Marine Evolutionary Biology, Zoological Institute, Christian-Albrechts-Universität Kiel, Am Botanischen Garten 1-9, 24118 Kiel, Germany; Evolutionary Ecology of Marine Fishes, Helmholtz-Centre for Ocean Research Kiel (GEOMAR), 24105 Kiel, Germany; Institute of Clinical Molecular Biology (IKMB), Christian-Albrechts-Universität Kiel, Rosalind-Franklin-Straße 12, 24105 Kiel, Germany

**Author notes:** Corresponding author: Freya Adele Pappert, Marine Evolutionary Biology, Zoological Institute, Christian-Albrechts-Universität Kiel, Am Botanischen Garten 1-9, 24118 Kiel, Germany.

**Keywords:** ageing, parental investment, sex role, pregnancy

## Abstract

Offspring quantity and quality are known to vary according to parental age, with most studies focusing on the mother’s age, who produces costly eggs and often carries out pregnancy, hampering to distinction between trans-generational age effects due to egg quality or physiological deterioration. We investigated the ramification of parental age on the offspring in the broad-nosed pipefish *Syngnathus typhle*, a fish species with male pregnancy, allowing us to separate these two female traits. By mating parents of different sizes we examined the impact of parental age on offspring number, size and gene expression. Our results show that older parents produced more and larger-sized offspring. However, we revealed intriguing insights into the differential gene expression patterns in offspring, strongly influenced by the paternal lineage but minimally affected by maternal age. Offspring from old fathers exhibited notable changes in gene expression profiles, particularly related to cell cycle regulation, metabolism, protein synthesis, stress response, DNA repair and neurogenesis. Our findings provide valuable insights into the role of pregnancy in shaping offspring physiology. Moreover, we recognize the value of assessing a broader range of species that have evolved with sex-specific differences in parental investment vs. gamete provisioning, as the age of either the mother or father may hold greater significance than the other in influencing offspring fitness.

## 1. Introduction

The burden of parental care and the energy investment necessary for reproduction are often on different scales between males and females. In many species, energy investment differs based on the size of the gametes (known as anisogamy), with eggs being typically much larger than sperm and therefore regarded as more costly to produce (1). This primary difference is supposedly at the root of sexual selection, with females having a higher cost in offspring investment compared to males and in turn being the choosy sex (2). Furthermore, we find that in numerous animals, females have also the burden of pregnancy, and in mammals frequently also lactation and child rearing. This is no small feat, as live-bearing, also known as viviparity, is considered one of the most significant forms of parental investment (3). This reproductive strategy enables the embryo to remain inside the womb until it has fully developed, providing protection from external threats, and ensuring the best possible conditions for embryogenesis. Viviparity has evolved independently more than 150 times in vertebrates, including fishes, amphibians, reptiles, and once in mammals (3).

Experimental studies on parental investment and brood quality have focused primarily on maternal characteristics, seeing them as a major component of the offspring’s own fitness, as females typically invest more not only in egg production but also in carrying and nurturing (4). Nonetheless, in the animal kingdom, designation of sex (egg and/or sperm producer) and their respective sex-role (i.e pregnancy, courting, etc.) can be more ambiguous (5). This is exemplified very nicely in teleost fish, that exhibit an immense variety of sexual systems from gonochorism to synchronous/sequential hermaphroditism (6,7). For example, the yellow-tail clownfish (*Amphiprion clarkii*) has a protandrous sex change, where the male will become a dominant female once the previous female dies provided he is physically the biggest (8), or the Okinawa rubble gobiid fish (*Trimma okinawae*) with cue sizes inducing a bi-directional sex change (9). The realm of bony fish have also seen the evolution of many varying forms of paternal care, from egg guarding, mouth brooding to varying degrees of pregnancy (10–12). Some of the most dedicated male parents can be accredited to the *Syngnathidae* family, which has evolved a range of male pregnant brooding forms from eggs carried on the males’ tails (e.g, *Nerophis ophidion*) to fully formed male pregnancies in brood pouches equipped with placenta-like system (i.e, *Hippocampus erectus*), providing the embryo with nutrients, oxygen and immunological components (13–16). Unfortunately, our understanding of how these diverse sex roles and associated life history trade-offs influence offspring fitness remains limited and gaining a deeper understanding could be crucial for several important areas in biology, such as evolutionary and reproductive biology, life history theory and even conservation strategies by helping manage populations with unique reproductive strategies. For this reason we used the broad-nosed pipefish *Syngnathus typhle* in this study, which not only has pregnant fathers who develop a brood pouch with a placenta-like structure housing the embryos (17), but also shows sex-role reversal with females competing for access to males during the breeding season (18,19). As their brood pouch is size-limited and the females produce on average more eggs than what the males can brood during the same period, males end up being the sex that is more reproductively constrained (20,21). The strength of sexual selection is reversed in this instance, with males being the choosier sex seeking out larger and more attractive females (19).

Our main aim is to understand how maternal and paternal size, used as a proxy for age, affects offspring quantity and quality in a sex-role reversed species with male pregnancy. This unique model species allows us to disentangle the effects of egg and pregnancy on the next generation, usually two female-specific traits, and investigate the value of the eggs provided by the mother and the role of the pregnant father. Across many taxa, parental age at the time of copulation affects the number of offspring produced, their quality and fitness, and sometimes their ultimate lifespan (22,23). With increasing age genetic mutations and epigenetic variations accumulate, telomere are shorter, the immune system weaker, altered cell communication leads to cancer, increasing the likelihood of having offspring with birth defects or decreased lifespan (24). In the common shag (*Phalacrocorax aristotelis*), an avian species, chicks produced by older parents experienced greater telomere loss during their growth period (25). Generally, older parents may have offspring with shorter lifespan due to increased frailty and/or more rapid ageing (22,25). The life experiences of parents can potentially influence the phenotype of their offspring through a process known as transgenerational plasticity (26,27). This mechanism is believed to facilitate the adaptation of embryos to their expected environment, potentially enhancing their overall fitness (26). This rapid acclimatisation primarily rely on epigenetic modifications, as opposed to genetic changes that require the accumulation of mutations over multiple generations (28,29). Over the course of a lifetime and influenced by lifestyle choices, epigenetic modifications can gradually accumulate due to both internal and external stressors. These modifications can disrupt cell communication and molecular processes, potentially affecting future generations as they are passed down to the embryo (30). Current understanding suggests that it is the biological age, reflecting physiological deterioration over time, rather than the chronological age in years lived, that plays a significant role in transmitting transgenerational effects on the health of offspring (23,24). Nonetheless, age-related decline is not universal, with some species experiencing rapid, negligible, or non-existing senescence (31,32).

Due to the previously mentioned differences in reproductive investment between males and females, trade-offs are to be considered when either parent decides to invest energy in their own offspring (33). Sex-specific effects of how much energy a father or a mother allocates into rearing their offspring is often in a fine balance with what the environment provides for nourishment and how physiologically fit an individual is (34). Studies in mice or in Asian elephants revealed that offspring from older mothers or even grandmothers had a shorter lifespan (35,36). Maternal ageing is hypothesised to affect the evolution of ageing, as it is her age, which appears to weigh more heavily on the mortality and/or survival of offspring (37,38), especially if in the particular species the female is predominantly responsible for pregnancy, breastfeeding, and rearing of the young (2,36). Markers measured in the mother such as oxidative stress or telomere length at birth, can affect the biological age of the offspring and such effects are strongest when resource availability is scarce (39). However, particularly in species where the young are not born precocial and are thus very vulnerable, having a young mother may lead to a higher offspring death (40). As she is physiologically and/or intellectually not mature enough to sustain a pregnancy or care adequately for her progeny, and may rather trade off her own somatic maintenance and survival against investing too much in her young (40). However, the effects of maternal age on offspring are also not always present; for example in invertebrates, such as the cockroaches, maternal age appeared to have no effect on brood quality (41). Equally, fertility does not necessarily always decline with age, with insects and fish often increasing offspring output with advancing age and increasing body size, providing additional nutrition (42,42,43). On the other hand, paternal age can also negatively affect the lifespan and health of offspring. Declining sperm quantity and quality, accumulation of germline mutations (23,44), and changes in resource allocation to ejaculate contribute to reduced fitness in male offspring (45,46). In some species having an older father has been shown to decrease the lifespan of the offspring and lead to more chronic illnesses (47,48). Although older fathers may exhibit genetic and molecular alterations due to ageing or environmental stressors, certain animal species show a preference for larger and older males (46,49), perceiving their longevity as an indicator of superior genes that could enhance offspring fitness, such as having fathers that will be stronger at protecting them from aggressors and/or provide higher quality paternal care, among other things (50,51). Mate choice plays a crucial role in the success of future generations, with reproductive trade-offs and species-specific senescence influencing the selection of the most suitable mate. Thus, to better understand the underlying reproductive trade-offs between sex and/or sex role and age effects on offspring fitness, it is important to consider the complexities and nuances of senescence and reproduction strategies between different species.

Thus our study wanted to answer how does the increased paternal investment (i.e., sex role) in *Syngnathus typhle* and the father’s age affect offspring, compared to the mother’s egg provisioning (i.e., sex)? Does the mother’s genetic background in the egg still count for more, compared to the father’s costly time and energy investment throughout pregnancy? Furthermore, is somatic age truly detrimental to both reproduction and offspring health in a species where fertility increases with age? To try and answer all our questions, we mated both old and young *S. typhle* in different combinations, i.e., old males with old females (OM x OF), old males with young females (OM x YF), young females with old males (YF x OM) and young females with young males (YM x YF). We measured quantitative parameters such as the duration of pregnancy, the total count of the offspring and their morphological measurements at birth. For the qualitative side, we opted to do full transcriptome mRNA sequencing (RNA-Seq) of at least five juveniles from each couple, to analyse what genes in the offspring may be differentially expressed based on the parental mating combination. We suggest that the effects of paternal age may surpass those of the mother, as the father’s energy investment in the offspring is more resource-intensive and is expected to vary based on his body size and life history. The findings of this study are important as they shed light on factors that could impact reproductive biology and offspring fitness in non-conventional sex-role species.

## 2. Methods

### 2.1 Sample collection

In this study, *Syngnathus typhle* was collected in late spring from two locations: the Limfjord in Trudsø, Denmark (56°47’N; 8°66’E) and the southwest Baltic Sea (54°39’N; 10°19’E). The specimens were brought in separately to the GEOMAR Institute where they were divided by sex and placed in 100-liter tanks at either 30 ppt (Limfjord) or 18 ppt (Baltic) salinity, maintained at a temperature of 14°C. To keep experimental conditions the same and to acclimate the pipefish collected from Trudsø to Baltic seawater, a slow filtered inlet drip was used over the course of a week till 18ppt was reached for all pipefish. During the quarantine period, the pipefish received two anti-parasite treatments with a combination of Malachitgrünoxalat (0.18g/100mL) and Formaldehyd (2.06g/100mL), applied on alternating days.

Before the experiment began, the body mass and length were measured to determine their age categories, as the larger the fish, the older it was estimated (52). One male and one female from the same population were then placed together in 30 individual tanks based on their size, where they were kept for a 2-week mating period. There was a higher number of pairs from Trudsø (23 pairs: 5 OMxOF, 6 OMxYF, 6 YMxOF, 6 YMxYF) compared to Falkenstein (7 pairs: 2 OMxOF, 1 OMxYF, 2 YMxOF, 2 YMxYF). The difference in the number of pipefish from each site was due to fewer fish being available at the Falckenstein site. Then the duration of male pregnancy was recorded. Once the offspring were born, five individuals from each parent were sampled and euthanized with an overdose of Tricaine mesylate (MS -222; 500 mg/l; Sigma-Aldrich), measured for total body length and weight. Then the whole body was immediately preserved in RNA later, as their organs were too small and translucent for individual dissection. The samples were stored at 4 °C for three days before being transferred to -20 °C for long-term storage.

### 2.2 RNA isolation and mRNA sequencing

To extract RNA from whole-body tissues, the RNeasy Mini Kit from Qiagen (Venlo, Netherlands) was used following the manufacturer’s instructions for a total of 98 samples, roughly five fries from five different parents and when pooled 25 for each parental age group (25 OMxOF, 23 OMxYF, 25 YMxOF, and 25 YMxYF). The quantity of extracted RNA was measured using a Peqlab NanoDrop ND-1000 spectral photometer (Erlangen, Germany) and the samples were stored at - 80°C. Library preparation was performed at BGI Tech Solutions in Hong Kong using the DNBSEQ Eukaryotic Strand-specific mRNA library, and then stranded mRNA sequencing was performed on a DNBseq platform (150bp reads, 25M clean paired-end reads per sample).

### 2.3 Statistical analysis

Statistical analysis was done in Rstudio v.4.2.2 (53). To analyse the collected data on body size, pregnancy duration, and offspring quantity, we initially assessed the normal distribution of the data and calculated mean and standard deviation. Next, we conducted separate one-way ANOVAs for length and weight of the adult pipefish that successfully mated and gave birth to determine how significantly different young and old parent pipefish (11 old males, 14 old females, 13 young males, 10 young females. The same test was done to assess any differences in pregnancy duration between various mating groups. Additionally, for the total offspring count and their morphological measurements at birth, we performed a one-way ANOVA test, followed by a Tukey’s Honest Significant Differences (HSD) test. The HSD test allowed us to conduct a closer pairwise comparison between the different groups. With regards to the RNA-Seq data, we performed a PERMANOVA (Permutational Multivariate Analysis of Variance Using Distance Matrices) using the “adonis2” function, “bray” method and family as strata. We did this analysis on logarithmically transformed count data to determine whether there were significant differences in transcript expression between parental mating combinations and their sampling location. After normalizing for differences in library size, filtering out low counts, and scaling to counts per million (cpm) as described in point 2.4, we performed a regularized log transformation procedure (rlog) on the data for a principal component analysis (PCA) to detect group clustering and possible outliers (Fig. S3), which were subsequently removed (Fig. 2A).

### 2.4 Transcriptome analysis

Quality control of the resulting reads from the Illumina sequencing was carried out using FastQC v.0.11.9 (54), and trimming was performed using Fastp v.0.20.1 (55). Subsequently, the reads were aligned to a whole genome assembly of *Syngnathus typhle* (BioProject ID: PRJNA947442) using STAR v.2.7.9a (56). Transcript abundance was calculated using TPMCalculator (57) in both raw read counts and transcripts per million (TPM). We used Orthofinder v.2.4.0 (58) to assign functional information to annotated genes with a species list including the following accessions: *Takifugu rubripes* (rubripes.fTakRub1.2), *Syngnathus rostellatus* (GCF_901709675.1, fSynAcu1.2), *Syngnathus acus* (GCF_901709675.1, GCF_901709675.1_fSynAcu1.2), *Nerophis ophidion* (unpublished), *Oryzias latipes* (ASM223467v1), *Hippocampus erectus* (PRJNA613176), *Hippocampus comes* (H_comes_QL1_v1, GCA_001891065.1), *Lepisosteus oculatus* (LepOcu1), *Gasterosteus aculeatus* (BROADS1), and *Danio rerio* (GRCz11). Differential gene expression analysis was performed on the RNA-Seq data obtained from illumina in Rstudio v.4.2.2 (53), using *S. acus* orthology annotations. The edgeR package v.3.40.2 (59) was used for scaling to counts per million (cpm) and normalising for differences in library size, and genes with less than 10 counts in at least five libraries were discarded. We then normalised for composition bias by trimmed mean of M values (TMM) with calcNormFactors function, which calculates a set of normalization factors for each sample thereby eliminating composition biases between libraries. After which we used the limma package v.3.54.1 (60), as a linear model-based method that uses empirical Bayes approach and voom method (converts count data to continuous scale) to identify DEGs. Limma is a software package useful in accounting for batch effects and mitigating other sources of technical variation, while voom has the best power of methods that controls the type I error rate and is specifically designed to handle library sizes that vary by an order of magnitude or more (61). This capability was particularly crucial in our experiment because we wanted to reduce any batch effects and type I errors in our differential gene expression results due to pooling five individuals per parent and thus not being able to entirely avoid direct parental effects. The differential expression analysis comparing several groups was done by creating a matrix of independent contrasts, this way it was possible to perform a one-way analysis of deviance (ANODEV) for each gene (i.e. makeContrast (OMxOFvsYMxYF = OMxOF - YMxYF, OMxOFvsOMxYF = OMxOF - OMxYF, …, levels = design). After which we estimated coefficients of interest between groups to have log fold changes with the function contrasts.fit and finally we performed an empirical Bayes moderation of the estimated variance which shrinks sample variance and applies a moderate t-test to identify DEGs. For further downstream analysis we also only used the resulting adjusted P values below 0.05 ensure that the observed treatment effects were real.

### 2.5 Orthology and enrichment analysis

We used a homology-based search of *Danio rerio* with OrthoFinder to annotate differentially expressed genes (DEGs)for gene ontology analysis. The *Danio rerio* orthologues of the differentially expressed genes per contrast group were then uploaded to g:Profiler (last accessed in March 2023) to conduct functional profiling based on the offspring’s parental mating age combination (OMxOF vs YMxYF, OMxOF vs YMxOF, OMxOF vs OMxYF, YMxYF vs YMxOF, YMxYF vs OMxYF). We set the statistical domain scope to “Only annotated genes” and the significance threshold to “g:SCS threshold” at 0.05. We also treated numeric IDs as “ENTREZGENE_ACC” and limited the data sources to GO biological processes and biological pathways from the Kyoto Encyclopaedia of Genes and Genomes (KEGG) and Reactome (REAC) databases. The results of the analysis are presented in Tables S1.

## 3. Results

### 3.1. Quantitative differences between offspring of different parental age combinations

Before beginning the experiment, the age of the parents was determined by their body size, with larger individuals being assumed to be older (52). A one-way ANOVA test revealed a significant difference in total body size between young and old females (P = 5.2e-12) and for weight (P = 3.69e-10), as well as young and old males (length, P = 4.6e-05; weight, P = 0.00413). Old females had a mean size of 19.36 cm (SD = 0.72), young females 14.56 cm (SD = 1.05), old males 17.05 cm (SD = 1.47) and young males 14.38 cm (SD = 1.11). Supplementary material Figures S1 A-D provide additional boxplot results of body weight and length of parents.

At the beginning of the experiment, we combined 23 pairs of male and female pipefish from Trudsø, combinations were 5 OM x OF, 6 OM x YF, 6 YM x OF and 6 YM x YF; and in further 7 mating combinations with Falckenstein pipefish with 2 OM x OF, 1 OM x YF, 2 YM x OF, 2 YM x YF. Of the original 30 *S. typhle* couples 24 successfully mated, with males accepting the eggs and following through with the pregnancy. We removed the five couples that did not successfully mate or where the males rejected the eggs from any further analysis. These included 1 OMxOF, 2 YMxYF, 1 OMxYF (Denmark population) and 1 YMxYF, 1 OMxYF (Germany population). From the 24 pregnant males, we tracked their pregnancy duration and found no significant difference between the four different groups of mating combinations (Fig. 1A). The average pregnancy time for each group was as follows: OMxOF 24.7 d (SD = 1.97), OMxYF 23.8 d (SD = 2.39), YmxOF 23.5 d (SD = 2.14), and YMxYF 25 d (SD = 1.58). Once the fry was born, they were counted and with a post-hoc Tukey’s HSD test we found the strongest significant difference (P = 0.0013) between OMxOF and OMxYF (Fig. 1B) and less so between YMxYF and OMxOF (P = 0.027). OM x OF had the highest number of offspring with an average of 68 (SD = 25.5), and OMxYF had the lowest average with 15.4 (SD = 9.1), while the other two groups were somewhere in between with YMxOF 46 (SD = 18.4) and YMxYF 31.4 (SD = 20.8). Yet, there was generally very high variability within families. From the pipefish juveniles (five per family) that we collected right at birth for gene expression sequencing, we also measured body length (Fig. 1C) and weight. We found no significant difference for weight but a significant difference for body length between the old parental combination (OMxOF) and all other groups (Fig. 1C).

**Fig. 1:**
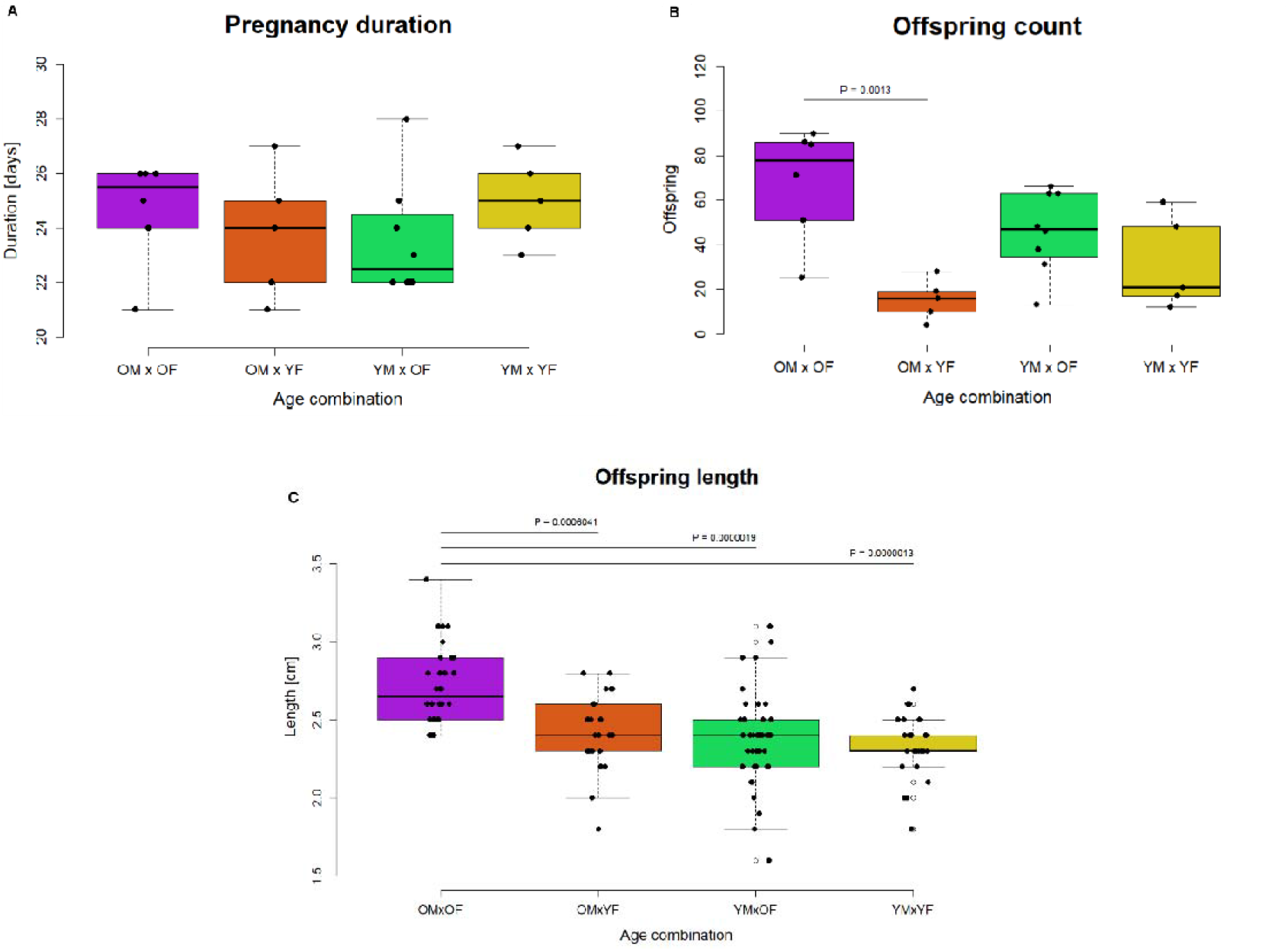
Boxplot representations of various quantitative measurements between different combinations of parental age of mating. In purple old males with old females, in orange old males with young females, in green young males with old females, and in yellow young males with young females. Highly significant P values from Tukey’s HSD test are marked above the boxplots with a line designating the comparison groups. A) Box plot of duration of the pregnancy in days. B) Boxplot of the offspring count per parental group. C) Boxplot of total body length immediately after birth of five offspring per parent in centimetres.

### 3.2. Age of father plays a critical role in offspring DEGs

After conducting a PERMANOVA, with factors being parental age combinations and location, on the 26072 transcripts from the sequenced *S. typhle* offspring, we found that parental age combinations were significant (P = 0.013), while the difference between sampling locations was not significant (P = 0.116), presumably as both parental populations were kept under the same conditions for several weeks before mating. However, there was some significant interaction between the combination of parental age and location (P = 0.009). Nonetheless, with a PCA we saw no differences between parental populations in four PCs (Fig. S2 A-C) and were thus confident that parental sampling location had no meaningful effect on the offspring and could group them together. When we then distinguished between the four different parental age groups we noticed that PC1 accounted for 20% of the variation, with the individual offspring clustering predominantly according to the father’s age (Fig. 1A). One can observe that on the left side of the plot individual offspring predominantly clustered based on parents from OMxOF and OMxYF combinations, thus old father being the common denominator, while on the right were offspring from YMxYF and YMxOF (Fig.1A), here young fathers being the common denominator.

The differential gene expression analysis produced distinct numbers of transcripts for offspring from different parental age combinations (Fig. 1B). When the age of both parents changed, 2874 genes were differentially expressed when comparing OMxOF vs. YMxYF, and 1716 for OMxYF vs. YMxOF (Fig.1B). To account for the age influence of both parents, 1417 genes were differentially expressed for OMxOF vs YMxOF, and 4630 for YMxYF vs OMxYF (Fig.1B). In contrast, changes in mother’s age produced the lowest number of DEGs, with 97 for OMxOF vs. OMxYF and 61 for YMxYF vs YMxOF (Fig.1B). We also analysed the DEGs for each comparative group to determine the number of genes that were upregulated or downregulated. Figure 2 from C to H, shows volcano plots of various log-fold change (logFC). The majority of the differentially expressed genes were between logFC of -0.5 and +0.5. When the age of the female changed, very few genes exceeded this threshold (Fig. 2D and G). In contrast, when the age of the father changed, many more genes had a higher logFC above 0.5 (Fig. 2E and F).

**Fig. 2:**
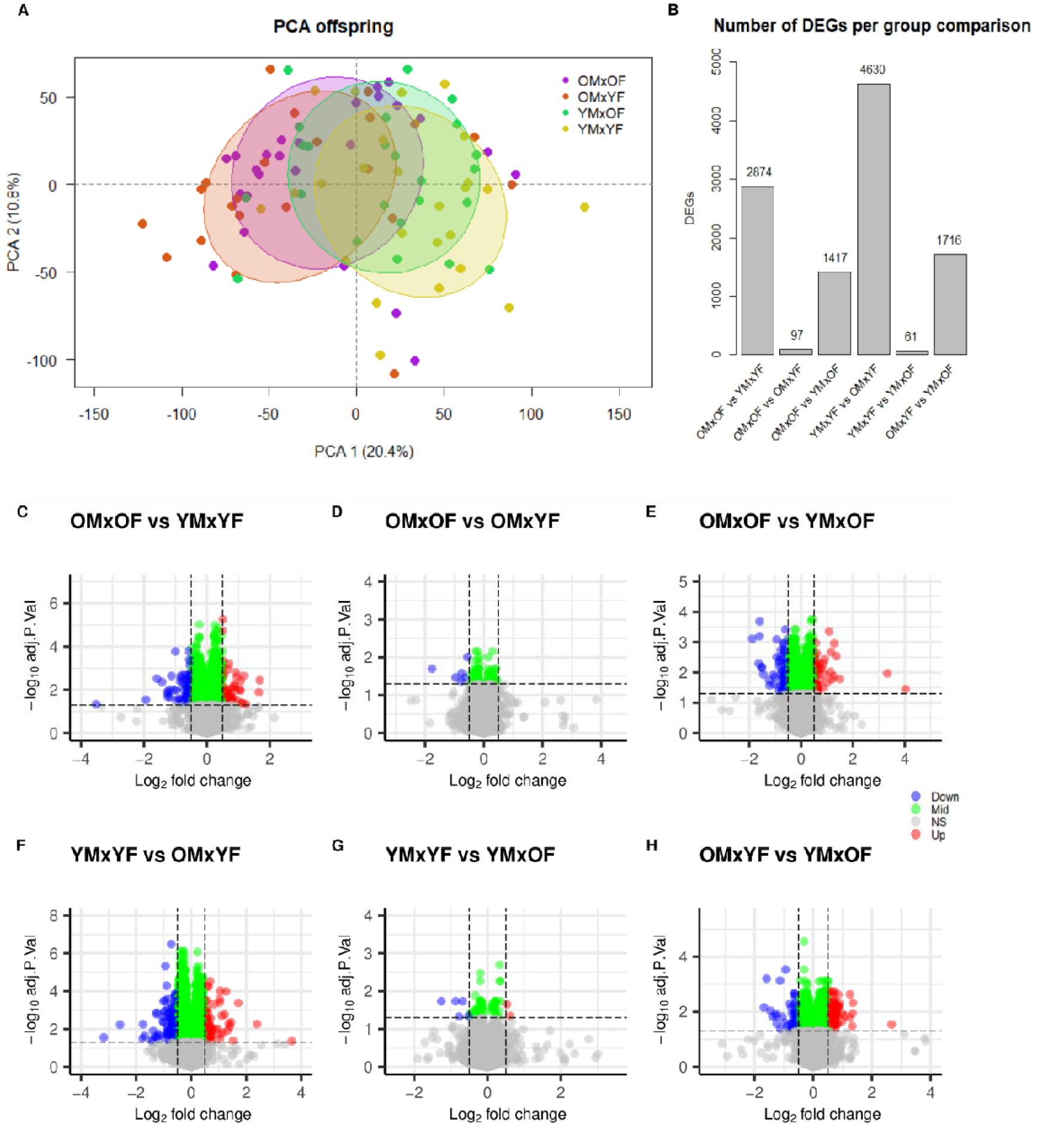
Results of the differential analysis of gene expression. A) Principal component analysis (PCA) of pipefish offspring grouped by parental combination: OMxOF (purple), OMxYF (brown), YMxOF (green), and YMxYF (yellow). B) Bar graph showing the number of differentially expressed genes (DEG) for each contrast comparison. C-H) Volcano plots showing the DEG analysis results for different contrast groups. The x-axis represents the log2 fold change, and the y-axis represents the negative log10 of the adjusted P-value. Grey dots represent non-significant values (adj.p.val > 0.05), red dots indicate genes with a logFC above 0.5 and adj.p.val < 0.05, blue dots indicate genes with a logFC below -0.5 and adj.p.val < 0.05, and green dots have adj.p.val < 0.05 with logFC between -0.05 and 0.05.

Results from the functional profiling of the DEGs in each comparison group using g:profiler, showed no enriched or over-represented categories for the OMxOF vs. OMxYF and YMxYF vs. YMxOF groups due to the low number of DEGs. On the contrary, when the age of the father changed, we found pathways enriched in categories such as cell cycle, cell stress and response, protein synthesis and regulation, signal transduction, metabolism and more (Table 1 and S2). Generally, the vast majority of overlapping pathways were found between OMxOF vs. YMxYF and YMxYF vs. OMxYF (Table 1). In the first group (OMxOF vs. YMxYF) it was complicated to disentangle what effects may be paternal or maternal, but we are confident that in the second group (YMxYF vs. OMxYF) effects observed are driven by male age. One of the few pathways that we found to be enriched in both OMxOF vs YMxOF and YMxYF vs OMxYF was the extension of telomeres, which activates telomere enzymes to add new telomere repeats onto the ends of chromosomes (62). Upon examining the genes involved in the extension of telomere pathway, we found that out of a total of 26 genes belonging to this pathway, 8 DEGs were present in OMxOF vs YMxOF and were all down-regulated in offspring from YMxOF, while for the other group there were 15 DEGs which were all upregulated for OMxYF (Tables S1 and S2). However, the majority of these genes were *replication factors* or *polymerase subunits*, and we found that some overlapped with other pathways such as DNA replication and DNA repair, as frequently these genes are involved in a complex network of molecular processes contributing to the maintenance of genomic integrity.

**Table 1:**
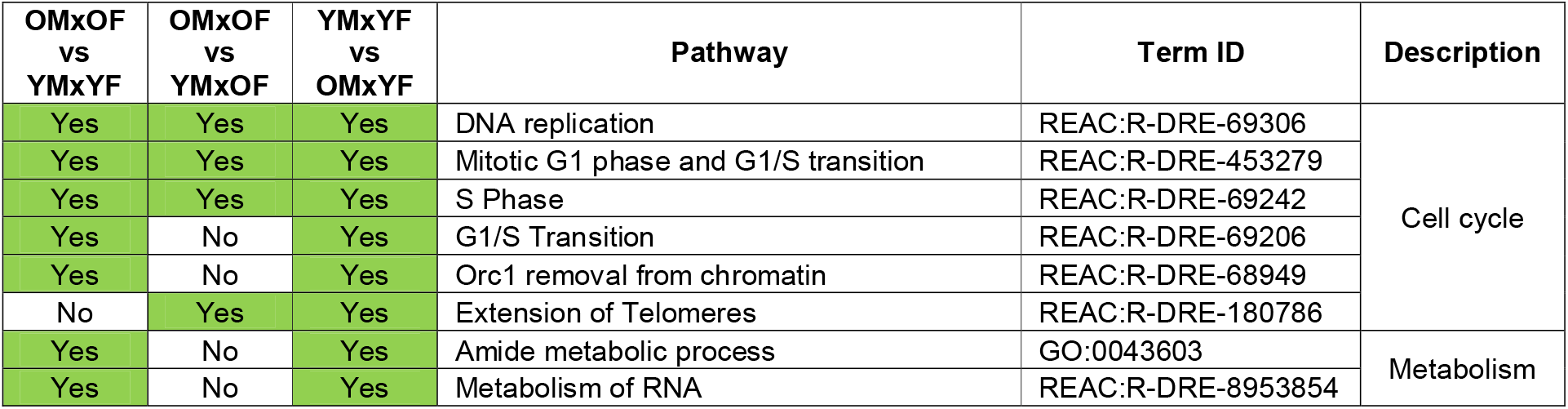
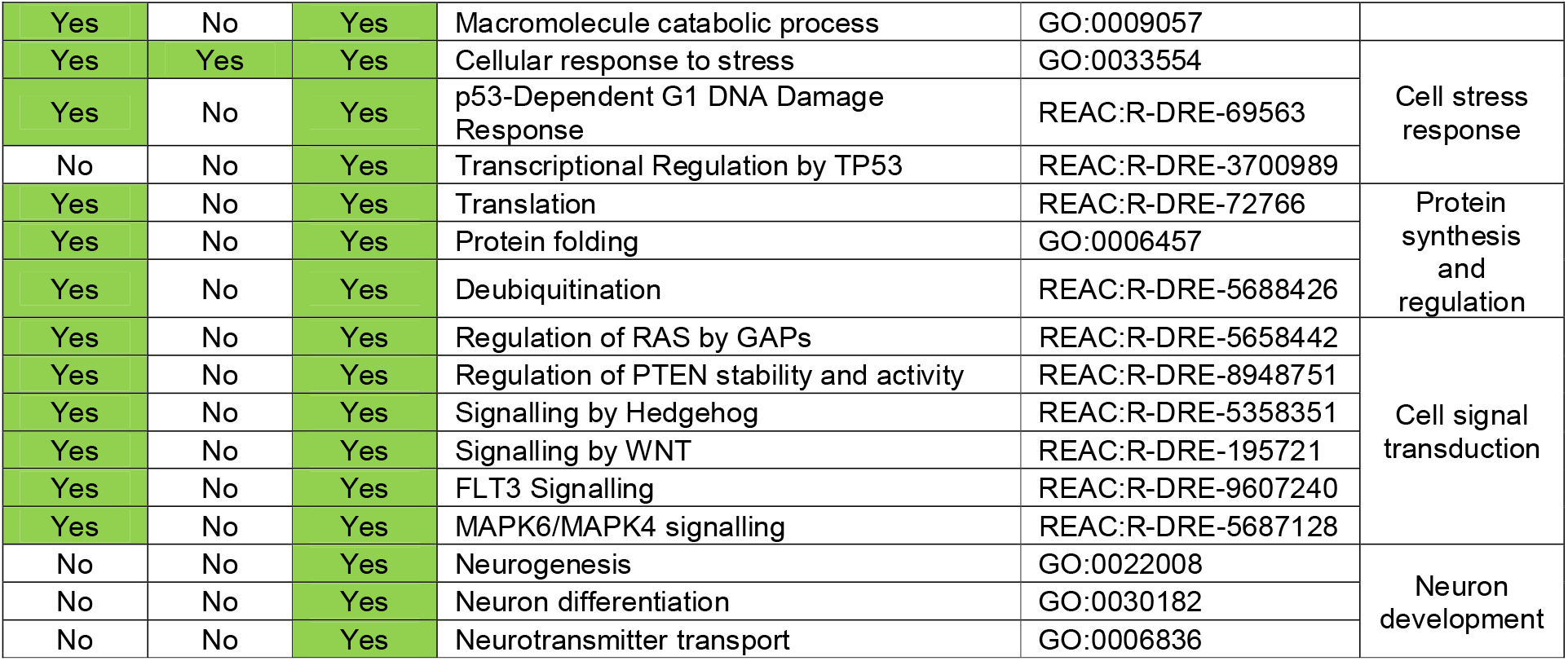
Summary table of pathways of interest from the gene set enrichment analysis. In the first three columns are the different parental group matches, where yes or no indicated if they had genes that were over-represented in a particular pathway.

There were numerous enriched pathways pertaining to metabolism, protein synthesis, cell signalling and transduction that occurred only when the offspring had an old father (Table 1 and S2). Over-represented genes in pathways such as translation, protein folding, metabolism of RNA, amide metabolic process, regulation of *PTEN* (Phosphatase and Tensin Homolog) stability and of *RAS* (Rat Sarcoma) by GAPs (GTPase-Activating Proteins), signalling by Hedgehog, and *MAPK* (Mitogen-activated protein kinase) signalling were predominantly if not exclusively up-regulated in offspring from older fathers (Tables S1 and S2). Although there were some overlaps in the genes involved in these pathways, many of them encoded for mitochondrial ribosomal proteins, heat shock proteins, cullin, chaperones, prefolding subunits, proteasome activators or subunits, splicing factors, and MAPK (Tables S1 and S2).

Furthermore, we found that the pathway for Orc1 removal from chromatin was overrepresented in OMxOF vs YMxYF and YMxYF vs. OMxYF, with approximately 27 and 43 genes, respectively. Notably, 25 of the 27 Orc1-related genes were upregulated in OMxOF, while all 43 genes were upregulated in OMxYF (Table S2). Orc1 is a crucial protein in the initiation of DNA replication in eukaryotic cells, and it has been found to have a strong impact on gene expression (63). On a similar note, offspring from old fathers had enriched p53 pathways, which are involved in the response to DNA damage and other cellular stressors. The p53 protein is a transcription factor that regulates the expression of genes involved in cell cycle arrest, DNA repair, apoptosis, and senescence (64). The majority of genes involved in this pathway were up-regulated in offspring with an older father, regardless of the mother’s age. These included replication factors, cyclin-dependent and checkpoint kinases, and tp53-regulating kinases (Table S1). Furthermore, *lamtor3-5* were upregulated, which are important components of the Ragulator-Rag complex, that is responsible for regulating cell metabolism, trafficking in eukaryotic cells and also the mTOR pathway as it responds to amino acid levels in the cell (65).

We found several GO terms for neurogenesis, neuron differentiation, neurotransmitter transport and so forth (Table 1 and S2), only for DEGs between YMxYF vs OMxYF. In particular, neurogenesis alone (pathway includes 900 genes) was associated with 200 DEGs, of which 177 were up-regulated in offspring from YMxYF (Tables S1 and S2).

## Discussion

We aimed to study how maternal and paternal age affects offspring quantity and quality in *Syngnathus typhle*, a sex-role reversed species with male pregnancy. We measured numerous variables, including body size and weight of the pipefish, pregnancy duration and number of offspring at birth. We observed major differences in the number of offspring at birth based on parental age (Fig.1B), despite male and female age not affecting pregnancy time or duration (Fig. 1A). On average, older parents had the highest number of offspring, which can likely be attributed to the larger brood pouch of male pipefish and the ability of larger females to produce more eggs. Interestingly, the lowest number of fry came from couples where the father was old and the mother was young, rather than from couples where both parents were young (Fig.1B). This finding suggests that older males may be less willing to accept eggs from younger females and may wait to fill their brood pouches to capacity when they encounter a larger and higher quality mate (66). Due to their polygamous mating behaviour, male *S. typhle* mate with multiple females, and embryos from different mothers can be found in a single pregnancy (67). We hypothesize that sexual selection, in this case male pipefish’s larger selection of partners, may explain the low number of offspring observed in parental couples with an old father and a young mother (OMxYF). We also observed a significant difference in body size measurements of the newly born juvenile pipefish between OMxOF and all other groups (Fig1.C). We know that egg size correlates positively with female body size in pipefish, so it is likely that having an older mother may provide a developmental advantage, as she will provide larger and more nutrient rich eggs, resulting in larger embryos (68). In the wild, bigger is generally better as the larger fry will be less vulnerable to predators, can swim faster for resources and are more competitive (69,70).

Our examination of transcriptome-wide differential gene expression revealed that the numerical disparity in transcripts among offspring from various combinations of parental age is primarily driven by the paternal lineage, with exceptionally many DEGs (4630) found when comparing offspring with both young parents (YMxYF) with those differing only in an older father (OMxYF) (Fig.2B). Interestingly, PCA also showed that juveniles grouped together according to their father’s age (Fig.2A), irrespective of the age of the mother. This suggests that paternal age and his experienced life history influences gene expression in *S. typhle* offspring. This may imply that the pregnant individual having longer contact with the developing embryos could influence epigenetic changes during embryogenesis and transmit non-genetic information, such as, immune history and microbial communities, impacting offspring phenotype and possibly influencing their survival and reproductive success in the future (17,26,27,71,72). Thus, the offspring overall profit from the increased size and age of mothers by starting off with a larger egg, but their gene expression at birth is likely more heavily influenced by the father. Research in species with female pregnancy have shown that mothers may have the stronger effect on offspring phenotype, as information passed from her ecological environment to the offspring is a more truthful signal (i.e salinity, temperature, food availability etc.) compared to the phenotype passed down from the father (73–75). Yet, in such cases, one cannot differentiate on whether it is due to the egg or truly the pregnancy, as the two are intermingled within the female. Our results suggest that in *S. typhle* the reverse is happening, as the developing embryos cannot predict their future environment, they will likely align their developmental trajectory with the paternal phenotype, thus making pregnancy a crucial element in transgenerational plasticity. Offspring differential gene expression being driven by paternal age is an indication that pregnancy, thus sex role and increased time investment in progeny, has a stronger effect on offspring characteristics compared to the maternal egg alone, and thus her genetic contribution. Unfortunately, from our RNAseq data we still cannot completely disentangle if these paternally inherited effects are predominantly epigenetically controlled and/or due to vertical transferral of immunological components or even due to the microbial pouch environment being transferred to the offspring, among other things.

Seeing as the paternal age appears to count for more in affecting offspring gene expression, are larger and thus older *S. typhle* fathers truly detrimental to offspring health? Functional profiling of DEGs revealed that the father’s age was mostly influencing cell cycle, stress response, protein synthesis and regulation, signal transduction, and metabolism of the offspring (Table 1). However, we found that all 43 DEGs included in the Orc1 removal from chromatin pathway were up-regulated in offspring with older fathers (Table S1 and S2), and as removal of Orc1 has been shown to cause defects in DNA replication and loss of integrity of chromatin structure (63), these changes can affect negatively gene expression and other cellular processes that depend on chromatin organization. Moreover, we also found many DEGs between YMxYF vs OMxYF that were involved in various cellular processes, such as protein synthesis and regulation, cell signal transduction, signalling by Hedgehog (Hh), MAPK6/MAPK4 (also known as ERK3/ERK4) signalling, regulation of *PTEN* stability, and regulation of *RAS* by GAPs, majority of which were upregulated offspring from OMxYF (Table 1 and Table S2). These pathways are primarily involved in growth and developmental processes and could indicate that age of the father at copulation is negatively influencing the rate of cell division in offspring, or possibly trying to mitigate DNA damage due to response to external stressors, which could both be linked back to the activation of telomerase (62,76,77). Unfortunately, limited research has been conducted specifically on the influence of parental age on these particular pathways. Studies have shown that de novo mutations in germ cells that occur at particularly higher rates in males, are then inherited and could subsequently lead to dysregulation of numerous cellular functions (78). There is also some evidence to suggest that with increasing age impairment of these pathways can occur and may lead to reduced fertility and developmental issues in new-borns (79,80), but so far there is no specific research of how these pathways are influencing offspring. The specific role on these pathways is nonetheless, for the most part, known. Hh signalling is particularly crucial in embryonic development, organogenesis and the renewal of various stem cell populations (81). The commonly denominated atypical MAP kinase ERK3/ERK4 pathway is lined to having both pro- and anti-oncogenic properties in various animal models (82,83). ERK/MAPK pathway is also known to play an important role in the regulation of neuronal survival and synaptic plasticity (84). This would go hand in hand with the enrichment of pathways involved in neurogenesis and neuron differentiation between YMxYF vs OMxYF (Table 1), in which 200 DEGs involved in the neurogenesis pathway alone had the vast majority (177 genes) upregulated in YMxYF (Table S1 and S2).

Nonetheless, we found that out of the 35 DEGs involved in the PTEN pathways between YMxYF vs OMxYF, there were 33 up-regulated in OMxYF (Table S2). *PTEN* encodes for a protein that plays a critical role in regulating cell growth and survival by inhibiting the PI3K/AKT signalling pathway (85), which in turn is crucial for regulating genetic stability and factors such as activating mTOR kinase inducing growth and translation, or by inhibiting transcription factors like FOXO family members thereby activating cell cycle progression and apoptosis (85). Both the activation of mTOR and inhibition of FOXO family members have a negative impact on health and longevity (86,87). Thus, PI3K/AKT is involved in driving tumour progression and up-regulation of *PTEN* leads to its suppression, so in this instance, offspring from young fathers are at a disadvantage compared to those from old fathers. Furthermore, we found that several downstream targets of TP53 were up-regulated in offspring with young fathers compared to offspring from old fathers, in particular two inducible tp53 proteins and two apoptosis-stimulating p53 proteins (Table S1). TP53 is a transcription factor that regulates the expression of a large number of genes involved in various cellular processes, including apoptosis, DNA repair, and cell cycle arrest (64). Studies have found a synergistic interaction between *PTEN* and p53, with the latter binding of to *PTEN* promoter regions and regulating their expression (85). Also the enriched pathway for regulation of *RAS* by GAPs, in YMxYF vs OMxYF, can function both as either a tumour suppressor or as an oncogene depending on the genetic variants within the gene, seeing as *RAS* genes were one of the first discovered oncogenes and their uncontrolled activation can lead to the development of tumours (88) and GAP can downregulate said *RAS* genes (89,90). It would appear in this instance that offspring from younger males may have a propensity for developing tumours and have an upregulated cell cycle activity. Unfortunately, we cannot confirm this with biopsies thus limiting our assumptions on possible negative health affects being passed on by either older or younger pregnant fathers. In the future we aim to follow up on the morphological development of the offspring and possible health deterioration or tumorigenesis, to then compare with our current results.

Overall, seeing as many genes involved in Orc1 removal from chromatin were upregulated in offspring from older fathers, leading to a potential increase DNA replication defects and overall loss of chromatin integrity, we think this would result in long-term negative health effects. Despite the fact that the offspring from older fathers also had an up-regulation of PTEN-related transcripts, possibly leading to an inhibition of the PI3K/AKT pathway and reducing the risk of tumorigenesis being ultimately beneficial. Particularly as there is growing body of literature showing that higher *PTEN* activity can influence neuronal survival and growth, contributing to various neurodegenerative disorders (91). As we also found several neuronal developmental pathways enriched for DEGs between offspring from YMxYF and those from OMxYF, with many transcripts being down-regulated in juvenile from old parents, we should also consider the possibility that the *PTEN* may be impacting negatively embryonic neurogenesis and further research is needed to confirm such effects. We propose that the significance of pregnancy in transgenerational plasticity in pipefish may be primarily attributed to the fact that older fathers have likely experienced a distinct life history compared to younger fathers, including potential exposure to pathogens or environmental stressors (i.e., nutrient availability, temperature fluctuations) which can then be modulating offspring phenotype.

Taken together, these findings suggest that theories about male and female senescence affecting offspring health should be reassessed as parental investment appears to be a more important driver than just egg versus sperm production. Moreover, we highlight the importance of moving beyond traditional animal models that oversimplify aging as a binary division between male and female reproductive systems. Instead, we must acknowledge the complexity of nature, where gender roles can be fluid and ageing can be influenced by trade-offs in life history. Studying reproductive biology in non-conventional sex-role species, helps us improve our understanding of the evolutionary trade-offs between male and female reproductive investment strategies, as well as their impact on transgenerational plasticity.

## Conclusion

This study focused on investigating the effects of maternal and paternal size, used as proxy for age, on offspring quantity and quality in *Syngnathus typhle*, a species with male pregnancy. While parental age did not affect pregnancy duration, older parents produced a higher quantity and larger sized offspring, likely due to larger male brood pouches and the ability of larger females to produce more and bigger eggs. Larger juveniles potentially have a higher survival rate in the wild, as they will be stronger and faster swimmers, giving them a advantages in food search and evading predators. However, the study also revealed that differential gene expression in offspring was strongly influenced by the paternal lineage, particularly in offspring from old fathers, and practically unaffected by maternal age. Seeing as *S. typhle* males are the pregnant sex, our results highlighting the importance of pregnancy in shaping offspring fitness. Offspring from couples with an old father exhibited differential gene expression profiles related to cell cycle regulation, metabolism, protein synthesis, stress response, DNA repair and neurogenesis. The upregulation of genes involved in telomere extension and chromatin organization indicated potential impacts on DNA replication integrity in offspring from old fathers. Nonetheless, the upregulation of PTEN-related transcripts in offspring from older fathers may offer protection against tumorigenesis. These findings indicate that the developing embryos are influenced by paternal age, but more research is needed to determine the long-term health effects. Our study ultimately stresses the importance of using animal models with varying forms of parental care and sex-roles, when studying the effects of parental age on offspring health and fitness, as such findings may possibly provide insight into non-genetic inheritance, developmental biology and reproductive medicine.

## Ethics statement

The work was carried out in accordance with the German animal welfare law and with the ethical approval given by the Schleswig-Holstein Ministerium für Energiewende, Landwirtschaft, Umwelt, Natur und Ditgitalisierung (MELUND) (permit no. 1288/2021). No wild endangered species were used in this investigation.

## Authors’ contributions

F.A.P and O.R. planned the project. F.A.P and D.K. carried out the experiment. F.A.P. analysed the data, interpreted the results and wrote the article. A.D. performed the orthology analysis and provided help with the GO profiling. O.R. helped during the field work and supervised the project. O.R. supported the writing of the manuscript.

## Acknowledgements

We gratefully acknowledge Mareike Marten for her valuable input and expertise as a lab technician and our animal-care technicians Fabian Wendt and Johannes Hasse for providing additional care and support with the animals.

## Funding

Financial support was provided by the Deutsche Forschungsgemeinschaft (DFG, German Research Foundation) through the Research Training Group for Translational Evolutionary Research (RTG 2501 TransEvo).

## Conflict of interest declaration

The authors declare no conflicts of interest.

## Data Access

Supplemental information include both additional figures (Fig. S1-S3), the *Danio rerio* orthologues of the significant differentially expressed genes in offspring pipefish between the various contrast groups (Table S1) and the gene set enrichment analysis results (Table S2). The raw sequencing data and metadata used in this study is available from the National Centre for Biotechnology Information (NCBI) Sequence Read Archive (SRA) under BioProject ID PRJNA973859 and accession number SUB13263866.

## Notes

https://submit.ncbi.nlm.nih.gov/subs/bioproject/SUB13263866/overview

### Competing Interest Statement

The authors have declared no competing interest.

