## Supplementary material for "The effect of parental age on the quantity and quality of offspring in *Syngnathus typhle*, a species with male pregnancy"

Supplementary materials

**Fig. S1**: Represented are boxplots of morphological measurement results of length and weight of pipefish parents. The x-axis represents the age groups, with old male (OM), young male (YM), old female (OF), and young female (YF). The y-axis represents the measured parameters. In A) and B), the length of the pipefish is represented in centimeters [cm] with panel A) depicting males and panel B) females, while in C) and D), the weight is represented in grams [g], with equally C) including males and D) including females. A) the mean total body size for OM was 17.05 cm (SD = 1.47) and for YM 14.38 cm (SD = 1.11), (OM vs. YM [cm] ANOVA, P = 4.6e-05). B) OF total body size average was 19.36 cm (SD = 0.72), and for YF 14.56 cm (SD = 1.05), (OF vs. YF [cm] ANOVA, P = 5.2e-12). C) OM had a mean body weight of 1.88 g (SD = 0.58) and YM of 1.27 g (SD = 0.36), (OM vs. YM [g] ANOVA, P = 0.00413). D) OF had an average weight of 3.28 g (SD = 0.41), and YF of 1.42 g (SD = 0.44), (OF vs. YF [g] ANOVA, P = 3.69e-10).

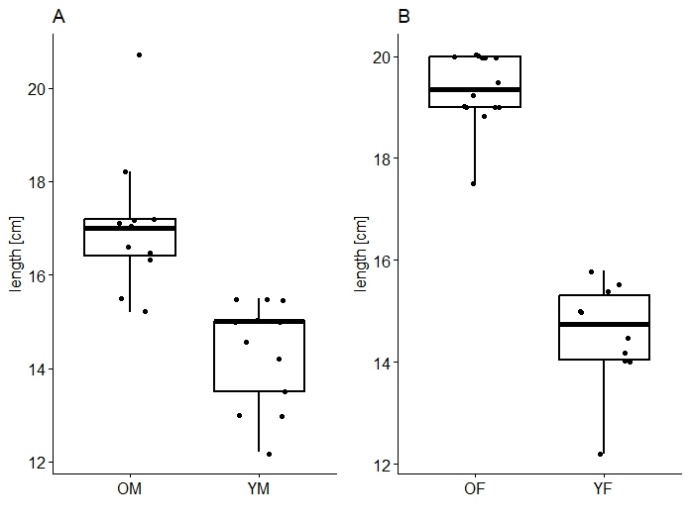

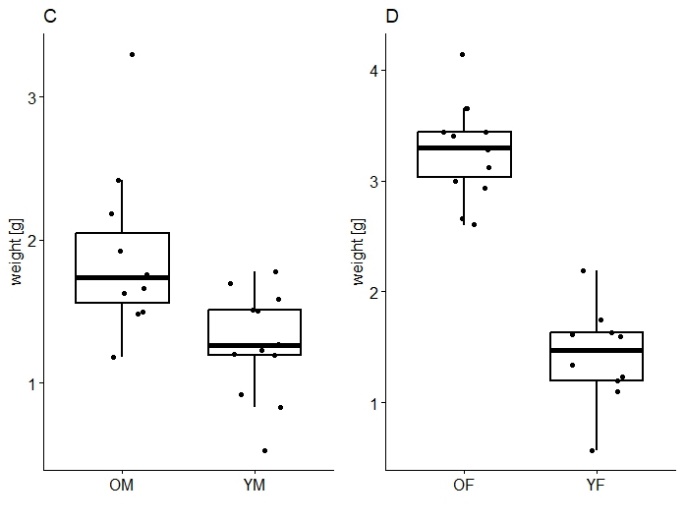

**Fig. S2 (A-C)**: PCA plots showing PC1 to PC4 on rlog transformed count data of all the offspring individuals based on if their parents came from Falckenstein (red) or Trudsø (blue). No obvious clustering was detected between offspring from different parental locations, which corroborated our PERMANOVA analysis (P > 0.05) that parental population didn’t have an effect.

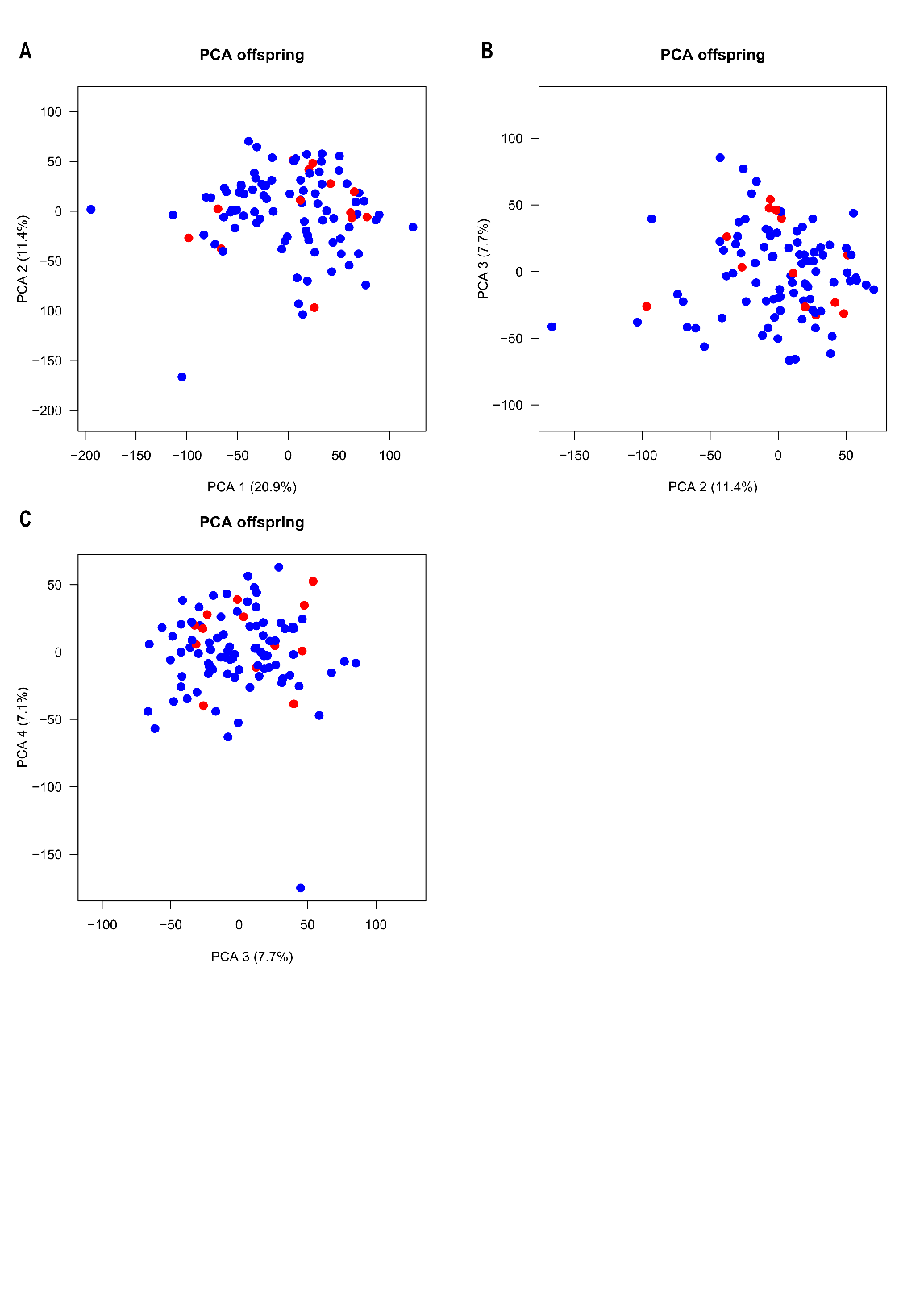

**Fig. S3**: The figure shows a principal component analysis (PCA) of rld transformed counts of pipefish offspring, grouped by parental combination: OMxOF (purple), OMxYF (brown), YMxOF (green), and YMxYF (yellow). PCA1 in all likelihood reflects the difference between offspring from old and young fathers. However, we detected two distinct outliers, one from OMxYF and one from YMxOF. These samples were distorting the distribution of the data and we decided to remove them for differential gene expression analysis to facilitate data normalization, enabling better analysis and modeling.

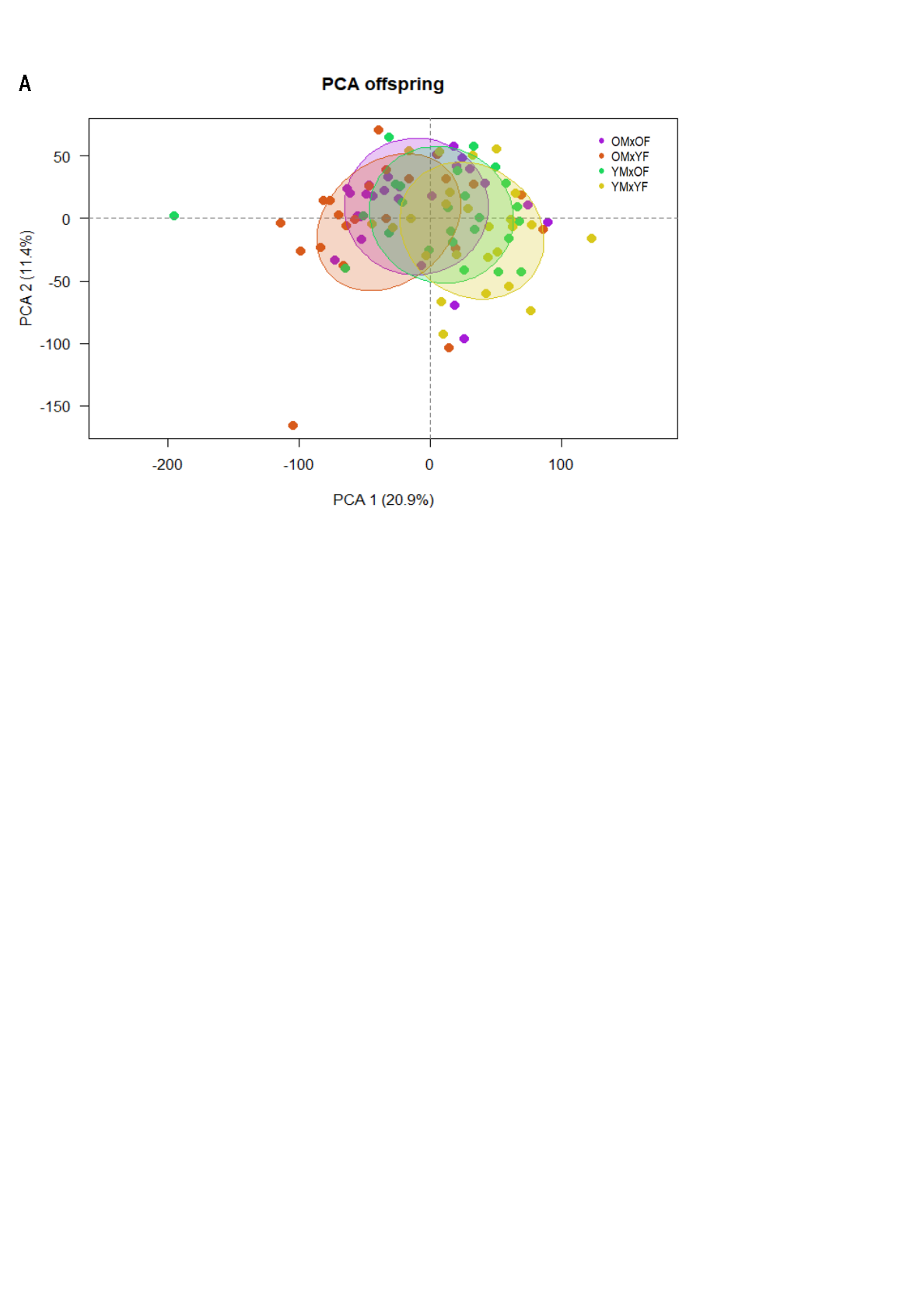
